# Genome-wide extraction of differentially methylated DNA regions using adapter-anchored proximity primers

**DOI:** 10.1101/2025.06.29.660377

**Authors:** Farzaneh Darbeheshti, Hayet Radia Zeggar, Hamzeh Salmani, Yibin Liu, Ruolin Liu, Viktor A. Adalsteinsson, G. Mike Makrigiorgos

## Abstract

The epigenetic deregulation of CpG islands (CGIs) plays a crucial role in cancer initiation and progression. CGIs comprise 1-2% of the human genome and are rich in differentially methylated regions (DMRs) that can serve as cancer biomarkers in clinical samples and liquid biopsies. Focusing epigenetic sequencing on CpG-rich sequences, including CGIs and avoiding non-informative regions, offers an efficient and sensitive approach for cancer identification and tracking, especially within samples containing excess of unaltered, normal DNA. To this end, we have developed Adaptor-anchored Methylation amplification via Proximity Primers (aMAPP), a versatile PCR-based enrichment method. aMAPP employs specially designed primers to selectively enrich either methylated or unmethylated CpGs, depending on the upstream methylation conversion method employed. aMAPP achieves high coverage of genome-wide CGIs and detects hundreds of DMRs in tumor samples compared to adjacent normal tissue using ultra-low depth sequencing (∼300,000 reads). It enables tracing of aberrant methylation down to allelic frequency 0.01% in dilutions of tumor DNA and in cell-free DNA samples, can be applied using picogram amounts of DNA, and can be adapted to enrich either small panels of cancer-specific DMRs, or the majority (>90%) of genomic CGIs and CpGs. aMAPP offers a simple, cost-effective, and highly sensitive approach for capturing the epigenetic footprint of genome-wide CpGs and identifying aberrantly methylated or un-methylated genomic regions.

**Graphical Abstract:** 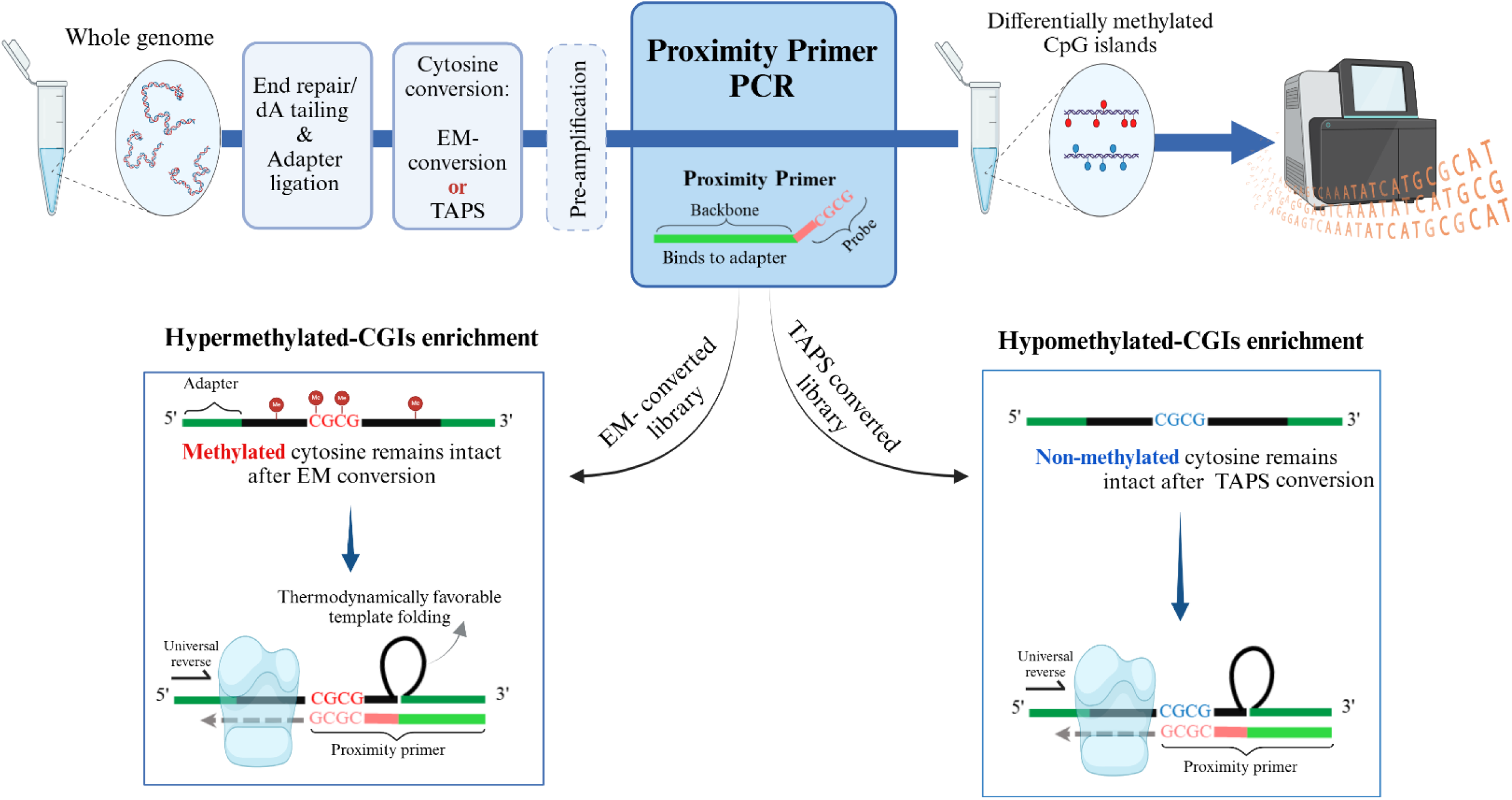

## Introduction

The significance of aberrant DNA methylation in cancer and other diseases has become increasingly clear over the past decade (1,2). DNA methylation markers are being used in cancer diagnosis, prognosis, and monitoring (3,4). Identification and tracking of differentially methylated DNA in cancer samples and liquid biopsies are commonly used as cancer biomarkers, similar to cancer-specific somatic mutations, but with added benefits such as tissue of origin (5–7). Discrete methylation changes in cancer often occur on CpG islands (CGIs), which comprise just 1-2% of the human genome (8). While whole-genome methylation profiling methods like whole genome bisulfite sequencing (WGBS) (9), enzymatic methyl-conversion sequencing (EM-seq) (10), and TET-assisted Pyridine Borane Sequencing (TAPS) (11) provide single-base resolution, they are inefficient for detecting aberrantly methylated CGIs in the presence of high excess of normal DNA. To enhance detection, methylation enrichment methods such as methylation immunoprecipitation (12) and methylation-specific restriction enzyme-based methods (13,14) have emerged, but they do not provide single-base resolution and have other limitations such as significant operational time and cost (15). Targeted methods employing cytosine deamination followed by allele-specific PCR, COLD-PCR, real time PCR, droplet PCR, isothermal amplification or hybridization capture using synthetic methylation probes (16–24) are limited in their genomic footprint while methylation array technologies are costly and not easily adaptable (25). Other techniques for enriching CG-rich regions, such as Reduced Representation Bisulfite Sequencing (RRBS) (26) and Heat-enriched Bisulfite Sequencing (27), methylation hybridization capture (28,29), and MCTA-Seq (30), have limited flexibility and cannot easily switch between targeting hyper-methylated or hypo-methylated regions at will.

Here, we describe a novel, simple, and cost-effective method, Adaptor-anchored Methylation Amplification using Proximity Primers (aMAPP) that enrich either hypermethylated or hypomethylated regions using a single workflow. We demonstrate that aMAPP surveys thousands of CGIs and CpG-rich regions using ultra-low sequencing depth and that the amplified regions are highly enriched in DMRs. This flexibility allows researchers to pivot from broad, pan-cancer screening to focused, cancer-specific panels using the same core workflow—a capability not easily achieved by other methods.

## Material and methods

### DNA samples and oligonucleotides

All ultramers, adapters, and primers were synthesized by Integrated DNA Technologies (IDT). The ultramers and adapters underwent polyacrylamide gel electrophoresis (PAGE) purification. Human fully-methylated and fully-nonmethylated control DNA was obtained from ZYMO Research (Catalog No. D5014). Five paired colorectal normal and tumor samples were sourced from Brigham and Women’s Hospital. All gDNA samples were sheared using Covaris ultrasonicator to achieve a mean fragment size of 150 bp and were stored in low TE buffer. Normal cfDNA samples from healthy volunteers were obtained from Brigham and Women’s Hospital, Boston, MA, with written informed consent.

### Library preparation

End Preparation and Ligation; It was performed using the NEBNext Ultra II DNA End Prep Kit for Illumina (Catalog No. E7645) according to the manufacturer’s protocol, with an input of 0.01-20 ng of DNA. The reaction was incubated at 20 °C for 30 minutes, followed by 65 °C for 30 minutes to deactivate the end prep. enzymes. Adaptor ligation was performed in the same reaction mix using the NEBNext Ultra II Ligation Module (Catalog No. E7645). This included 2.5 μl of 1.5 μM customized Y-shaped adapter, inspired by the TruSeq Illumina adapter, along with 1 μl of ligation enhancer and 30 μl of ligation master mix. For doing enzymatic conversion in the next step, we used methylated adapters in which all cytosines are methylated (Supplementary Table 1). The reaction was incubated at 20 °C for 30 minutes. Following adapter ligation, purification was carried out using AMPure XP beads at a 1.2X bead-to-sample ratio, with final elution in 28 μl of low TE buffer.

### Cytosine Conversion

The ligated DNA underwent either enzymatic methyl-conversion (EM-seq) or TET-assisted pyridine borane sequencing (TAPS). The former was performed using the NEBNext Methyl-seq Conversion Module (Catalog No. E7125S) according to the manufacturer’s protocol. The EM-seq protocol consists of two main steps: TET2 oxidation and APOBEC deamination. The converted DNA was then cleaned using 1.2X AMPure XP beads and eluted in 15 μl of nuclease-free water.

TAPS conversion was performed following published protocols (11). The ligated cfDNA was subsequently incubated in a 50-μL reaction mixture containing 50 mM Tris-HCl (pH=8.0), 5 mM α-ketoglutarate, 2 mM adenosine triphosphate, 2 mM dithiothreitol, 5 mM ascorbic acid, 40 μM ammonium iron (II) sulfate, 0.1% Tween-20, and TET2 enzyme (Catalog No. EM301-02, Vazyme) for 1 hour at 30 °C. Following incubation, a stop reagent was added to terminate the reaction, and the mixture was further incubated for 30 minutes at 37 °C. Reaction products were then purified using 1.8× AMPure XP beads. Oxidized DNA was then reduced in a 50 μl reaction containing 100 mM sodium acetate solution (pH=4.0) and 100 mM pic-borane (Alfa Aesar) for 6 hours at 37 °C and 850 rpm in the Eppendorf ThermoMixer. The product was purified by Zymo-IC column with Oligo Binding Buffer (Catalog No. D4060, ZYMO Research). For both EM-Seq and TAPS-converted libraries, pre-amplification was performed using NEBNext Q5U Master Mix (Catalog No. M0597) with primers targeting the adapters (Supplementary Table 1) for 8 cycles, following the manufacturer’s protocol.

### Proximity Primer-PCR (PP-PCR)

Methylation-specific amplification was carried out using novel and customized proximity primers composed of an anchor (binding to the adapter) and a probe (binding to CpG dinucleotides somewhere upstream of the adapter). These primers identify methylated cytosines that remain as cytosine after EM-seq and/or unmethylated cytosines that remain intact after TAPS, allowing DNA polymerase to initiate amplification (Supplementary Table 1). The proximity primers were used along with a universal primer, allowing them to act as forward and/or reverse primers depending on whether the probe target is on the top or bottom strand, or on both (Fig. 1d). PP-PCR reaction is done in a total volume of 50 μl, including 20 μl converted DNA from the previous step, 25 μl NEBNext Ultra II Q5 Master Mix, 5 μl of 10 μM proximity primer and universal reverse primer. The reaction is performed in an Eppendorf PCR machine with the following program: 98 °C for 30 seconds, followed by 25 cycles of 98 °C for 10 seconds, 65 °C for 75 seconds, and final extension at 65 °C for 5 minutes. After Proximity Primer-PCR and purification with 1,2X AMPure XP beads, the samples were ready for sequencing using Illumina platforms, as they possess partial Illumina adapters.

**Figure 1.**
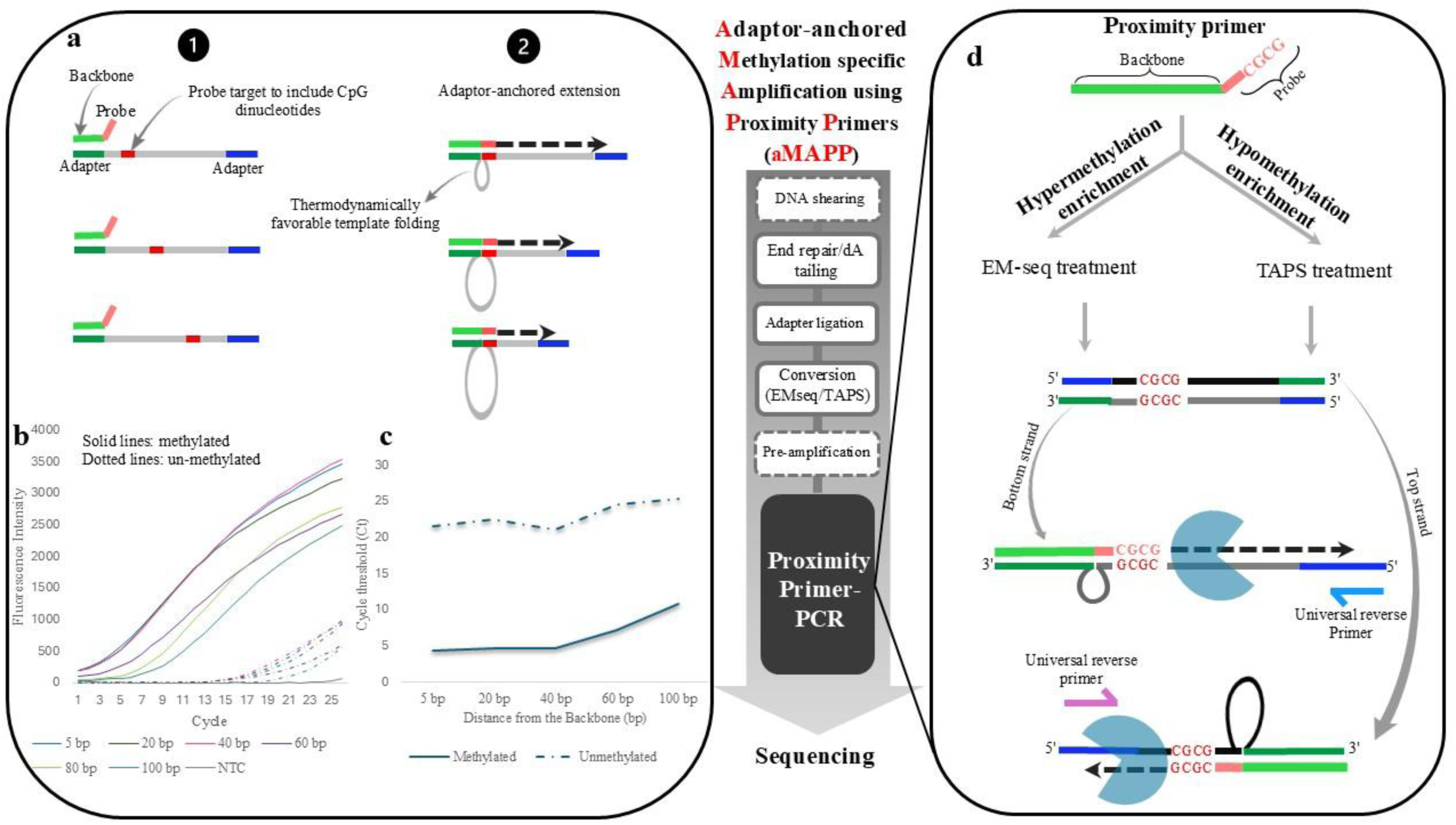
Overview of Methylation-Specific Amplification using Proximity Primers (MAPP). **a)** Schematic representation of the proximity primer-PCR concept. Proximity primers consist of two components: “Backbone” and “Probe”. The backbone functions as a universal primer complementary to the adapter sequence, while the probe (6-12 bp) is variable and targets a nearby sequence. The DNA strand can fold, allowing the probe to bind to its fully matched target, resulting to a deletion in the folded region. **b)** Demonstration of Proximity Primer-PCR using a proximity primer with a 9-bp probe (CGACCCGCG) to amplify different fragments from a single target, representing cfDNA or randomly fragmented gDNA. Converted methylated ultramers are depicted with solid lines and converted unmethylated ultramers with dashed lines. The probe targets are located at varying distances from the adapter. **c)** Plot illustrating the efficiency of Proximity Primer-PCR as a function of the distance between the probe target and the adapter. **d)** Application of Proximity Primer-PCR for methylation-specific amplification, showcasing a novel approach for extracting and enriching methylated, or un-methylated, CpG-containing genomic regions. The probe can bind to both the top and bottom strands, depending on target availability.

### Sequencing

Libraries were sequenced on either a MiSeq platform (2×250 bp, ∼50,000 reads/sample) or a NovaSeq X Plus platform (2×150 bp, ∼300 million reads/sample) by Genewiz Azenta Life Science.

### Data processing

#### Trimming and alignment

The raw sequencing reads were processed using Trim Galore to remove low-quality bases, adapters, and probe sequences. This step ensures that only high-quality sequences are retained for downstream analysis, reducing potential errors in alignment and methylation calling. The trimmed reads were aligned to a reference genome using either Bismark (31) or BWA-MEM (32), depending on the conversion method. After alignment, duplicate reads were identified and removed using Bismark’s deduplication tool. This step is crucial to ensure that each unique read is represented only once in downstream analyses, preventing bias in methylation statistics.

#### Extraction of methylation information

For EM-seq converted libraries, the methylation status of the aligned reads was extracted from the deduplicated BAM files using Bismark’s methylation extractor (31). While for TAPS converted libraries, methylation information is extracted using MethylDackel (33). To investigate the relationship between methylation status and CpG islands using Bedtools intersect (34), we intersected the deduplicated BAM file with a predefined set of CpG islands.

#### DMR analysis

To identify DMRs following aMAPP in paired tumor and normal tissue samples, we developed a pipeline to meet the following criteria: 1) Only the first 20 bases of the trimmed reads downstream of the proximity primer probe is analyzed. This condition ensures that we only include CpGs that are near the probe targets, which are expected to include 3 CpGs in their sequence. At least two additional CpGs must be present within the first 20 bp of the trimmed reads to be considered a potential DMR. 2) For sequences present in the tumor sample but absent in the normal samples, if the tumor sample paired-end coverage is ≥ four, those regions are considered unmethylated in the normal sample for EM-seq converted, while they are considered methylated for TAPS converted libraries. Otherwise, those regions are excluded from the analysis. 3) If coverage is present in both the tumor and normal samples, these are analyzed regardless of the coverage level. After data sorting based on these criteria, the DSS (Dispersion Shrinkage for Sequencing data) algorithm (35) is used for differential methylation analysis with the following settings: mean p-value = 0.05; minlen = 5; delta = 0.1; dis.merge = 300; minCG = 2. For analyzing DMRs in the standard libraries, the default DSS settings were applied across the entire read length.

## Results

### Proof of concept

Proximity primers consist of two components: the ‘backbone’ and the ‘probe.’ The backbone is complementary to an adapter sequence ligated to the interrogated DNA using a standard library preparation kit, while the probe (6–12 bp) contains CpGs and is complementary to sequences that potentially exist downstream on the same DNA strand. During the annealing stage of proximity primer-PCR (PP-PCR), the backbone anneals to the ligated adapter, then the DNA strand folds, enabling the 3’ end of the probe to bind a nearby matched target sequence, followed by primer extension (Fig. 1a). Similarly designed ‘loop primers’ or ‘super-selective’ primers were previously described for sensitive genotyping and PCR (36,37). To enable such primers to work with converted, randomly fragmented DNA (e.g. cfDNA) for methylation-specific enrichment of CpG-rich target sequences, we match the backbone to the adapter sequence ligated on every DNA strand. We first examined whether such primers could amplify proximal CpG-containing targets on the same DNA strand, at distances anywhere between 5-100 bp from the adapter. Long oligonucleotides (ultramers) mimicking de-amination converted methylated, or unmethylated, promoter regions of the RASSF1 gene were synthesized. These ultramers included the sequence of adapter sequences at each end, with probe targets positioned at gradually increasing distances from the adapter where the backbone of the primer is expected to bind first (Supplementary Table 2). Using a proximity primer with a 9-bp Probe (CGACCCGCG), we demonstrated via real time PCR that the proximity primer amplifies efficiently methylated ultramers when the probe target is located anywhere between 5 and 100 bp from the adapter sequence. In contrast, amplification of unmethylated ultramers was approximately 2¹⁵ (or 32,768 times) less efficient due to the absence of a suitable probe-binding site following de-amination and conversion of CpG to TG (Fig. 1b and 1c). Moreover, Sanger sequencing and Bioanalyzer fragment size assessment confirmed that PCR products contained the partial sequence deletions (Supplementary Fig. 1). These deletions are expected to occur following the thermodynamically favorable loop-folding when the targeted sequence is located at varying distances from the adapter.

Next, building on this concept, we developed a genome-wide enrichment method for capturing hypermethylated and/or hypomethylated CpG sites on genomic DNA. Depending on the conversion method selected, either the unmethylated CpGs (bisulfite, EM-Seq) or the methylated CpGs (TAPS) can be selectively converted (11). In the former case, using EM-Seq, proximity primers bind to adaptors and amplify methylated targets matching the probes. In the latter case, using TAPS, the same proximity primers bind to adaptors and amplify un-methylated targets with almost no change in the workflow. Hence, proximity primers would be expected to selectively amplify DNA fragments containing methylated CpGs that remain intact following enzymatic conversion or, alternatively, unmethylated CpGs that persist after TAPS (Fig. 1d).

To assess the ability of aMAPP in capturing methylated and unmethylated CpGs, we applied the method to fragmented gDNA from the NA12878 cell line converted using either EM-seq or TAPS as conversion methods. In both experiments, the proximity primer contained the probe sequence GACCCGCG, and a single protocol was used for enrichment. Following ultra-low depth sequencing, about 300,000 sequencing reads per sample, the CpG methylation status of the sequenced CpGs was determined. In parallel, we also conducted the same experiment using standard EM-Seq or TAPS libraries at the same sequencing depth. Standard EM-seq and TAPS libraries use the adapter sequence for amplification following the conversion step. Hence, there is no enrichment taking place, essentially representing ‘shallow’ whole genome methylation sequencing. Following aMAPP on EM-seq converted DNA, the density plots of methylation percentages for CpGs with paired-end depths ≥5 demonstrate strong enrichment for methylated CpGs, whereas aMAPP application on TAPS-converted DNA shows enrichment for unmethylated CpGs (Fig. 2a). In the EM-seq–converted library, using aMAPP, 88.8% of covered CpGs exhibit over 70% methylation. In contrast, in the TAPS-converted library, 96.8% of covered CpGs show less than 20% methylation. The data indicate that aMAPP extracts efficiently either methylated CpGs or un-methylated CpGs, depending on the conversion method applied. Worth noting, sequencing of standard EM-seq and TAPS libraries (coverage cutoff ≥2) shows a broad distribution of CpG methylation typical of whole genome methylation sequencing, with standard EM-seq showing 28.7% of covered CpGs having ≥70% methylation while standard TAPS yields 43.6% of covered CpGs having ≤20% methylation (Fig. 2a).

**Figure 2.**
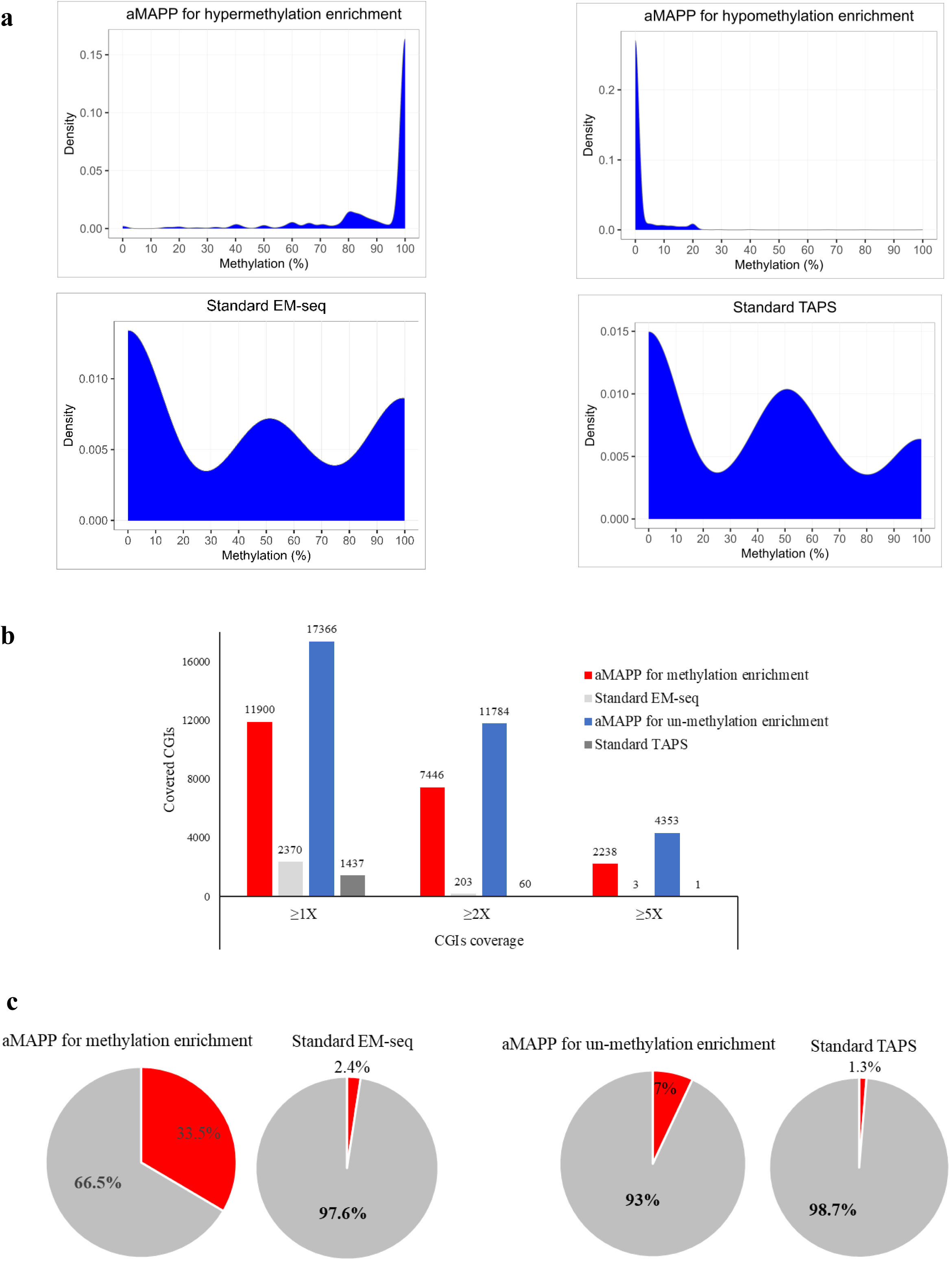
aMAPP efficiently captures both methylated and unmethylated CpG Islands (CGIs) using ultra-low depth sequencing. Methylation enrichment via aMAPP was performed on EM-seq-converted DNA, while un-methylation enrichment was conducted on TAPS-converted DNA. **a)** CpG methylation percentage and distribution in aMAPP libraries compared to standard libraries using sheared NA12878 gDNA. aMAPP selectively enriches methylated and unmethylated CpGs depending on the conversion method applied. **b)** Number of distinct CGIs covered using ultra-low depth sequencing in aMAPP libraries compared to standard control libraries of methylated and unmethylated control DNA. **c)** Ratio of CGI-associated reads to total reads in aMAPP libraries versus standard control libraries of methylated and unmethylated control DNA.

To assess the reproducibility of aMAPP, we performed a triplicate using the sheared gDNA from NA12878 cell line followed by EM-Seq and employing proximity primers containing the octamer probe GACCCGCG at the 3’end. Following ultra-low depth sequencing, we assessed the methylation status of the extracted CpGs. The methylation status of CpGs exhibited a strong positive correlation among the triplicates, with Pearson correlation coefficients of 0.88 and 0.91 for paired-end depths of ≥5 and ≥10, respectively (Supplementary Fig. 2a). We also examined triplicate samples of a colorectal tumor gDNA using the same protocol. Notably, 98% of CGIs with paired-end depths ≥5 in Replica-1 were also present in Replica-2 and 3 (Supplementary Fig. 2b), indicating the reproducibility of the proximity primer amplifications.

### aMAPP efficiently enriches either methylated or unmethylated CpGs from CpG islands using ultra-low depth sequencing

Next, we examined the aMAPP amplification footprint across genome-wide CpG islands (CGIs) using ultra-low depth sequencing. To this end, we applied aMAPP to human control methylated and unmethylated DNA for methylation and unmethylation enrichment, respectively, using again proximity primers with the octamer probe sequence GACCCGCG. We compared the efficiency of aMAPP in extracting CpGs from CGIs compared to standard EM-seq and TAPS libraries at the same sequencing depth.

In methylation enrichment, aMAPP yielded 36.6-fold and 746-fold more CGIs with paired-end depths of ≥2 and ≥5, respectively, compared to standard EM-seq. For un-methylation enrichment, aMAPP yielded 196.4-fold and 4353-fold more CGIs at paired-end depths of ≥2 and ≥5, respectively, compared to standard TAPS (Fig. 2b). Additionally, aMAPP significantly increases the proportion of CGI-related reads in library pools. Specifically, when applied to EM-seq converted DNA, the CGI reads vs. total reads ratio increased from 2.4% in standard EM-seq library to 33.5% in aMAPP library (Fig. 2c). This enrichment was less pronounced in the TAPS-converted library, where the CGI ratio increased from 1.3% to 7%.

### aMAPP performance with low DNA input and flexible targeting

We also assessed the influence of starting DNA input on the capability of aMAPP in extracting CpGs from CGIs, we examined its performance using decreasing amounts of input DNA. As a challenging, clinically relevant test, we employed cfDNA from healthy volunteers using inputs of 5ng, 1ng, 0.1ng, and 0.01ng. Applying aMAPP to EM-seq converted cfDNA using ultra-low-depth sequencing, ∼300,000 reads, resulted in extracting CpGs from 3,841 distinct CGIs with a 5ng input. The number of CGIs detected declines with DNA input. However, even at the lowest input level of 0.01ng, aMAPP successfully captured 905 CGIs, of which 258 overlapped with those detected in the 5ng sample (Supplementary Fig. 3).

By design, aMAPP offers enhanced flexibility for efficiently capturing cancer-specific CGIs by modifying probe sequences, particularly at the 3’ nucleotides. To evaluate this, various probes at the 3’ end of proximity primers were tested on control methylated DNA to assess their effectiveness in capturing methylated CGIs within the top 20 hyper-methylated CGIs for a given cancer, using ultra-low depth sequencing. The probe CGAACGCGAA successfully captured all 20 of the most common hypermethylated CGIs in colorectal cancer, whereas CGAACGCGAC captured 14 colorectal cancer-specific hypermethylated CGIs (Supplementary Fig. 4a). Interestingly, the latter probe demonstrated superior efficiency in capturing breast cancer-specific hypermethylated CGIs, covering 18 out of the 20 (Supplementary Fig. 4b). Notably, aMAPP consistently outperforms standard EM-seq in capturing cancer-specific hypermethylated CGIs using ultra-low depth sequencing, regardless of the probe used (Supplementary Fig. 4).

To assess aMAPP footprint modulation depending on applied conditions and the ability to capture CpGs throughout the genome, we applied aMAPP using a simultaneous combination of 3 proximity primers with different probes (CGCGAACGCGTA, CGAACGCGAA, AACGCG) on methylated control DNA that underwent EM-seq conversion. This experiment revealed that aMAPP could capture approximately 6.6 x 10^5^ distinct CpGs using ultra-low-depth sequencing. When the same experiment was repeated at moderate sequencing depth ∼10 million sequencing reads, the number of distinct CpGs covered increased to 10.7 million, which amounts to ∼37% of the 28.8 million of total CpGs genome-wide (38). Considering that ∼13 million genomic CpGs are contained in repetitive elements (38) that have lower success in alignment relative to non-repetitive sequences, it is likely that under the conditions applied, aMAPP surveys even more than 37% of non-repetitive element CpGs.

### aMAPP identifies differentially methylated regions (DMRs) in cancer samples using ultra-low depth sequencing

In the next step, the ability of aMAPP to detect DMRs in cancer tissue was evaluated using ultra-low depth sequencing. To achieve this, aMAPP was applied to both EM-seq- and TAPS-converted DNA from a colorectal tumor sample and its adjacent normal tissue. For both libraries, a single aMAPP protocol incorporating the same proximity primer was utilized.

The analysis of aMAPP on the EM-seq-converted libraries identified 293 hypermethylated DMRs, 158 of which exhibited a beta value ≥ 0.8 (Supplementary Table 3). These included well-known biomarkers for colorectal cancer, such as SEPTIN9 (39–41), GATA4 (42–44), CDH4 (45,46), FGF14 (47,48), IKZF1 (49), NKX6-2 (50), SLC6A2 (51), SOX1-OT (52), PRKCB (53), FOXB1 (54), MEIS1 (54), and EDIL3 (54), Fig. 3a. On the other hand, applying aMAPP on the TAPS-converted libraries revealed 55 hypomethylated DMRs. 29 of which showed a beta value ≤ -0.8 (Supplementary Table 4). These hypomethylated DMRs include well-known oncogenes previously reported as upregulated in colon cancer, including TWIST1 (55–60), PITX2 (61–63), FAM83D (64–66), CDC25A (67,68), ARID3A (69), as well as in other types of cancer like GNA12 (70–72) (Fig. 3a). In contrast, standard EM-seq and standard TAPS libraries at the same sequencing depth, ∼300×10^3^ reads, identified only 16 hypermethylated and 3 hypomethylated DMRs respectively, 84% of which had paired-end depth of one (Fig. 3b). Notably, by applying aMAPP, only 15.8% of the 348 detected DMRs had paired-end depth of one, while the remaining DMRs were supported by two or more read pairs.

**Figure 3.**
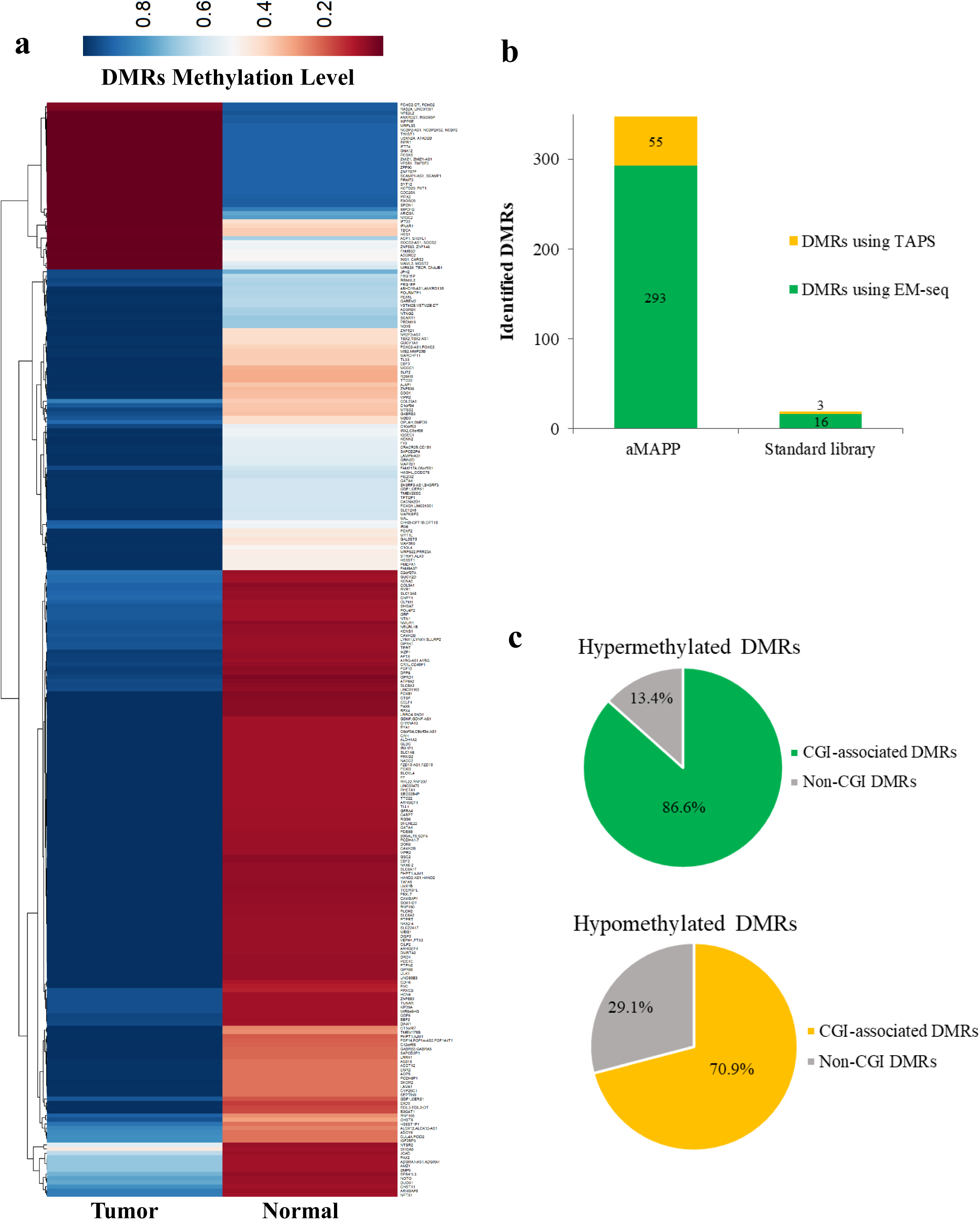
Detection of differentially methylated regions (DMRs) using aMAPP in a colorectal tumor sample through ultra-low depth sequencing, ∼300×10^3^ reads. **a)** Heatmap illustrating the methylation levels of detected DMRs, which annotated close a gene, in tumor tissue versus adjacent normal tissue. **b)** Total number of DMRs identified using aMAPP compared to standard library methods under ultra-low depth sequencing conditions. aMAPP applied to EM-seq-converted DNA detected 293 hypermethylated DMRs, and aMAPP applied to TAPS-converted DNA identified 55 hypomethylated DMRs. While standard EM-seq and standard TAPS libraries detected 16 and 3 DMRs, respectively. **c)** Proportion of DMRs associated with CpG Islands (CGIs), detected using aMAPP on EM-seq-converted DNA (hypermethylated DMRs) and aMAPP on TAPS-converted DNA (hypomethylated DMRs).

Among the DMRs detected using aMAPP, 86.6% of hypermethylated DMRs from EM-seq-converted DNA and 70.9% of hypomethylated DMRs from TAPS-converted DNA were associated with CGI regions (Fig. 3c).

### aMAPP enables detection of DMRs at low frequencies (0.01%) in both solid tumor and circulating-DNA samples using ultra-low-depth sequencing

To evaluate aMAPP’s ability to trace low-level methylation and non-methylation, we first applied it to sheared control methylated and unmethylated gDNA. For methylation tracing, methylated control DNA was diluted in unmethylated control DNA at ratios of 10% and 1%. Conversely, for non-methylation tracing, unmethylated control DNA was diluted in methylated control DNA at the same ratios. These two groups of DNA inputs were then processed using enzymatic conversion and TAPS, respectively. Our results demonstrate the strong enrichment effect of aMAPP. When applied to an EM-seq–converted library, aMAPP showed 70% methylation in a sample that was only 1% methylated. Conversely, when applied to a TAPS-converted library, it detected 51% non-methylation in a sample that was only 1% unmethylated. Interestingly, even the fully unmethylated (EM-seq–converted) and fully methylated (TAPS-converted) control samples showed 36% methylation and 26% non-methylation, respectively, when analyzed with aMAPP. This result probably reflects the fact that the commercially supplied control DNA is not fully 100% methylated or unmethylated, as the manufacturer states a purity of over 95%. Another reason may be the incomplete conversion of EM-seq (usually >99% conversion on C) and TAPS (>85% on 5mC). Overall, our data indicate that aMAPP effectively enhances the detection of low-level methylation and non-methylation, even in commercially available control methylated and unmethylated DNA samples (Supplementary Fig. 5).

To further assess the sensitivity of methylation detection, we tested aMAPP on samples designed to mimic extremely low levels of tumor DNA. This was applied by diluting the colorectal tumor DNA within its adjacent normal tissue and in pooled cfDNA from 5 healthy volunteers, followed by enzymatic EM-seq conversion. The data show detection of several tumor DNA targets down to 0.01% level. For example, Figure 4 illustrates one of the highly methylated CGIs located near the NKX6-2 gene (chr10:132783854-132789145; CpG: 642) which was identified in DMR analyses of the tumor and its adjacent normal tissue. When testing aMAPP on 0.01% spike-in tumor DNA within adjacent normal tissue, the methylated CGI tumor was detected, revealing the presence of tumor DNA (Fig. 4a). Similarly, in the 0.01% spike-in tumor DNA within normal cfDNA, aMAPP successfully captured methylated tumor DNA (Fig. 4b). Notably, normal cfDNA exhibited completely unmethylated CpGs in the same region, and no amplification was observed in adjacent normal tissue, confirming that this CGI is unmethylated in normal samples.

**Figure 4.**
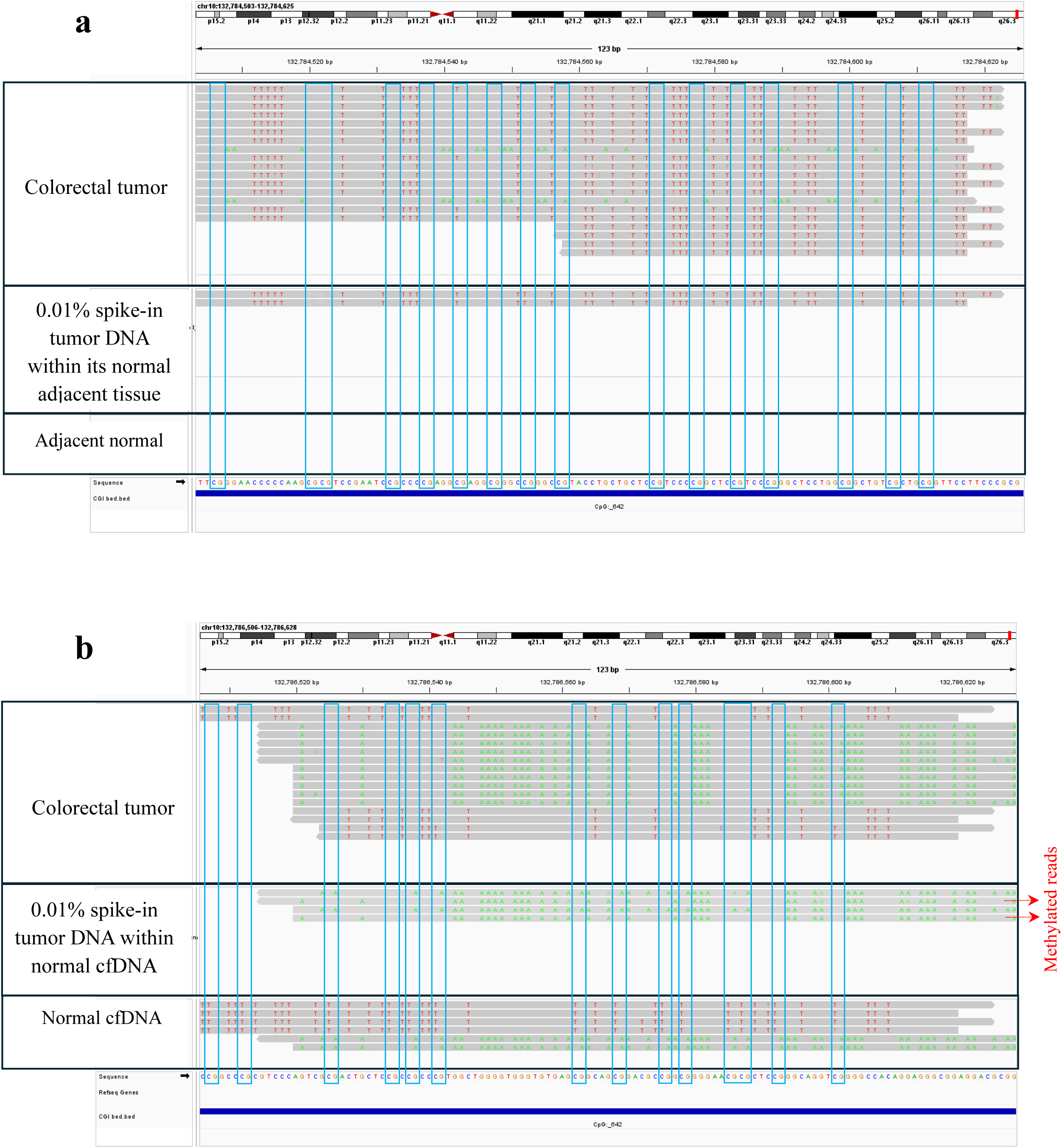
aMAPP detects DMRs at a 0.01% frequency in both solid tumor (a) and cfDNA (b) samples using ultra-low-depth sequencing, 3×10^3^ reads. Integrated Genome Viewer, IGV visualization of a highly methylated CGI (chr10:132783854-132789145; CpG: 642) identified in DMR analyses of colon tumor and adjacent normal tissue (referenced in Fig. 3a). This CGI is located near the NKX6-2 gene. **a)** In a 0.01% spike-in tumor DNA sample within adjacent normal tissue, aMAPP successfully detected methylated tumor DNA. No amplification was observed in normal tissue, confirming the unmethylated state of this CGI in normal samples. **b)** In a 0.01% spike-in tumor DNA sample within normal cfDNA, aMAPP effectively captured methylated tumor DNA in 2 out of 4 sequence reads. The same region in normal cfDNA showed completely unmethylated CpGs.

### DMRs detection in clinical samples using aMAPP at moderate sequencing depth

Although aMAPP enables detection of DMRs with ultra-low depth sequencing, ∼300,000 reads, even at low allelic frequencies, a more comprehensive DMR detection can be achieved using moderate sequencing depth. We thus applied aMAPP followed by ∼10 million reads on five colorectal tumors and their normal adjacent tissues, which underwent enzymatic EM-seq conversion. aMAPP detected 6664, 6691, 8834, 5669, and 4534 DMRs in tumors 1 to 5, respectively, of which 79.7%, 61.2%, 78.3%, 77.3%, and 74.1% fall within CGIs (Fig. 5a and Supplementary Table 5). Next, we assessed which CGIs across all five samples contain common DMRs. A total of 387 CGIs with DMRs were shared across all five samples, encompassing several well-characterized hypermethylated colorectal cancer biomarkers, including SEPTIN9, GATA4, CDH1, SFRP1, and SDC2, among others (Fig. 5b and Supplementary Table 6).

**Figure 5.**
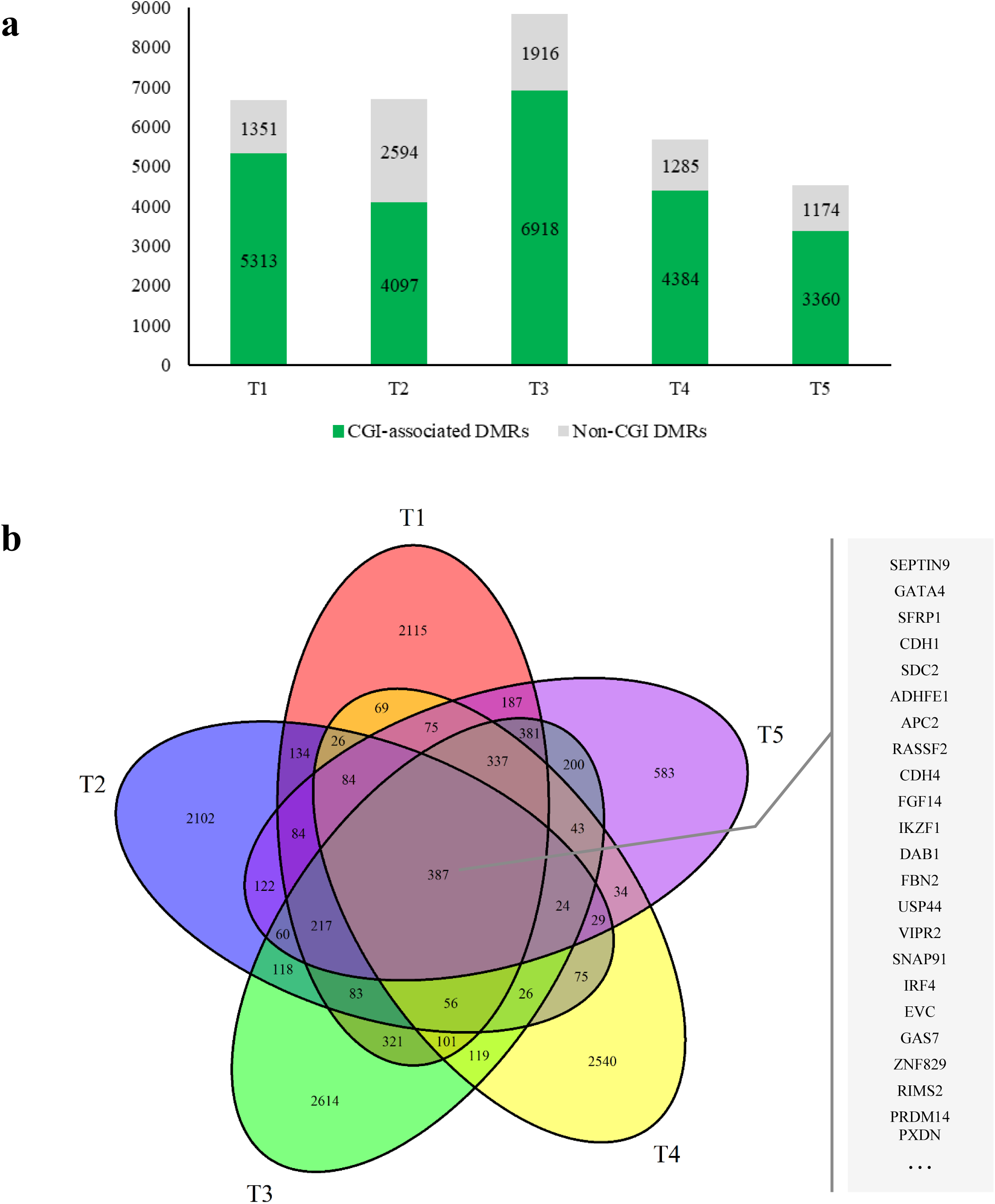
Broader DMR detection using aMAPP (probe sequence GACCCGCG) on EM-seq converted libraries followed by moderate sequencing depth (10M reads). **a)** Number of detected DMRs in five colorectal tumors compared to their normal adjacent tissues. **b)** Venn diagram illustrating common differentially methylated CpG Islands (DM-CGIs) among the five colorectal tumors. The common DM-CGIs shared across all five samples are associated with several hypermethylated colorectal biomarkers located in gene promoters, as listed.

## Discussion

Methylation detection is an established biomarker for cancer detection and monitoring, as well as in a broader horizon beyond oncology (73,74). We have developed aMAPP, a novel PCR-based enrichment method for the facile detection of hypermethylated and hypomethylated DMRs. Though prior techniques for genome-wide methylation profiling and enrichment have been developed (9,10,12,75), aMAPP is able to extract DMRs from CGIs and CpG-rich regions at single-base resolution for either hypermethylation or hypomethylation using a single protocol, and employing ultra-low depth sequencing. While aberrant hypermethylation in cancer is usually localized, predominantly occurring in CGIs and gene promoters, hypomethylation has a dual mechanism of action: loss of methylation in specific CGIs, including oncogenes, alongside global hypomethylation across the genome, including repetitive elements, which contributes to genomic instability (76–80). For example, LINE-1 hypomethylation in cfDNA has been proposed as a potential biomarker for early cancer detection and prognosis (81–84). Accordingly, adapting proximity primer probes in aMAPP could specifically target and assess hypomethylated DMRs on LINE-1. As another possibility, the methylation profiles of 200–250 DMRs enable tissue of origin prediction for about 20 different types of cancer (70,71). aMAPP proximity primers could potentially be adjusted to capture these tissue- or tumor-specific regions, providing a simple, sensitive, and cost-effective approach for detecting tissue of origin. By applying multiple proximity primers simultaneously and moderate-depth sequencing, one can capture almost all CGIs in the genome and extract biologically rich information. Accordingly, the ability to modulate the targets and footprint of the enriched genomic regions by adjusting the aMAPP proximity primer, following standard library preparation, offers practical advantages that make aMAPP amenable to multiple potential applications.

With regards to detecting tumor circulating DNA (ctDNA) with potential uses in early detection, tumor monitoring or detection of minimal residual disease, aMAPP can identify DMRs down to 0.01% allelic frequencies (∼100ppm) using ultra-low sequencing. Tumor informed approaches based on mutation tracking, and using duplex sequencing principles (85–87) offer a 10-100 times lower limit of detection, 1-10 ppm (88,89). However these rely on the availability of primary tumor tissue for analysis, which is not always possible to obtain (90). Merging mutation and methylation information in a single library preparation (91,92) may combine advantages of both tumor informed and tumor agnostic approaches. While aMAPP detection sensitivity of rare alleles is still not at the ppm-level, it is possible that methylation DMR information obtained via aMAPP could enable tissue-agnostic ctDNA detection and potentially supplement tumor informed approaches by providing tissue-of-origin information.

Longer probes or modulation of PCR cycling conditions using proximity primers can provide a more targeted approach, and inclusion of probe modifications, such as locked nucleic acid (LNA) incorporation, can potentially increase target specificity (93). While the current aMAPP application was demonstrated using EM-seq and TAPS conversion protocols, the technique would be expected to remain essentially unchanged using bisulfite or enzymatic approaches that achieve single-step methylation conversion (94). aMAPP operates with a single PCR step following standard library preparation, thus offering an accessible and cost-effective protocol, which provides speed and practical advantages over hybrid capture methylation panels (23).

In conclusion, aMAPP provides unique versatility, flexibility and efficiency in detecting epigenetic changes and identifying low-abundance DMRs, with potential for broad application in cancer and beyond.

## Acknowledgements

This study used the NA12878 cell line sample from the NINDS Repository and HCT116 DKO cell lines provided by ZYMO Research. The graphical abstract was created in BioRender. Darbeheshti, F. (2025) https://BioRender.com/swr9uc6.

## Author Contributions

G.M.M., F.D., and H.R.Z. jointly designed the study and engineered optimal protocols. F.D. and H.R.Z. performed the experiments. H.S. analyzed the aMAPP sequencing data. F.D. designed the adapter, wrote the first manuscript draft, and created the graphical abstract. H.S., F.D., and H.R.Z. contributed to the graphs. Y.L. contributed the TAPS protocol and discussed study implications. V.A.A. and R.L. interpreted the data, discussed study implications and contributed to the study design. G.M.M. proposed the use of proximity primers and supervised the study. All authors read, edited, and approved of the manuscript.

