## Supplementary Fig. for "Genome-wide extraction of differentially methylated DNA regions using adapter-anchored proximity primers"

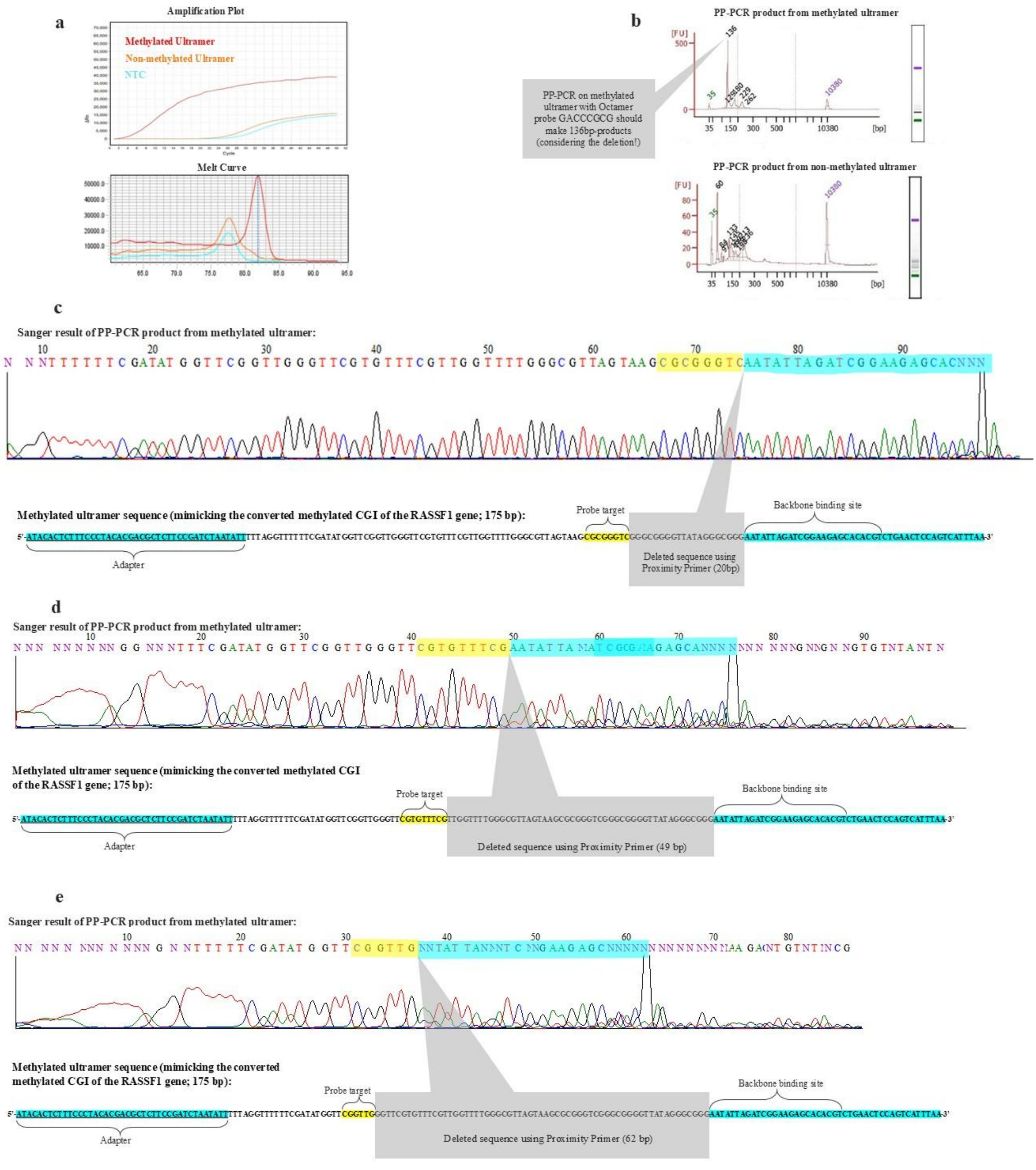


**Supplementary Fig. 1:** Confirmation experiments for Proximity Primer-PCR (PP-PCR) functionality.

**a)** Amplification and melt curve analyses related to PP-PCR using the octamer probe GACCCGCG on ultramers that mimic converted methylated and non-methylated CGI regions of the RASSF1 gene. **b)** Bioanalyzer results using high-sensitivity DNA chips to assess the fragment size of PP-PCR products. **c)** Sanger sequencing results of the PP-PCR product from methylated ultramers, confirming a 20-bp deletion sequence occurring in the folded region (the sequence between the adapter and probe target). **d)** Sanger sequencing results of the PP-PCR product using a nonamer probe CGAAACACG on methylated ultramers, confirming a 49-bp deletion sequence occurring in the folded region. **e)** Sanger sequencing results of the PP-PCR product using a hexamer probe CAACCG on methylated ultramers, confirming a 62-bp deletion sequence occurring in the folded region.

**a**

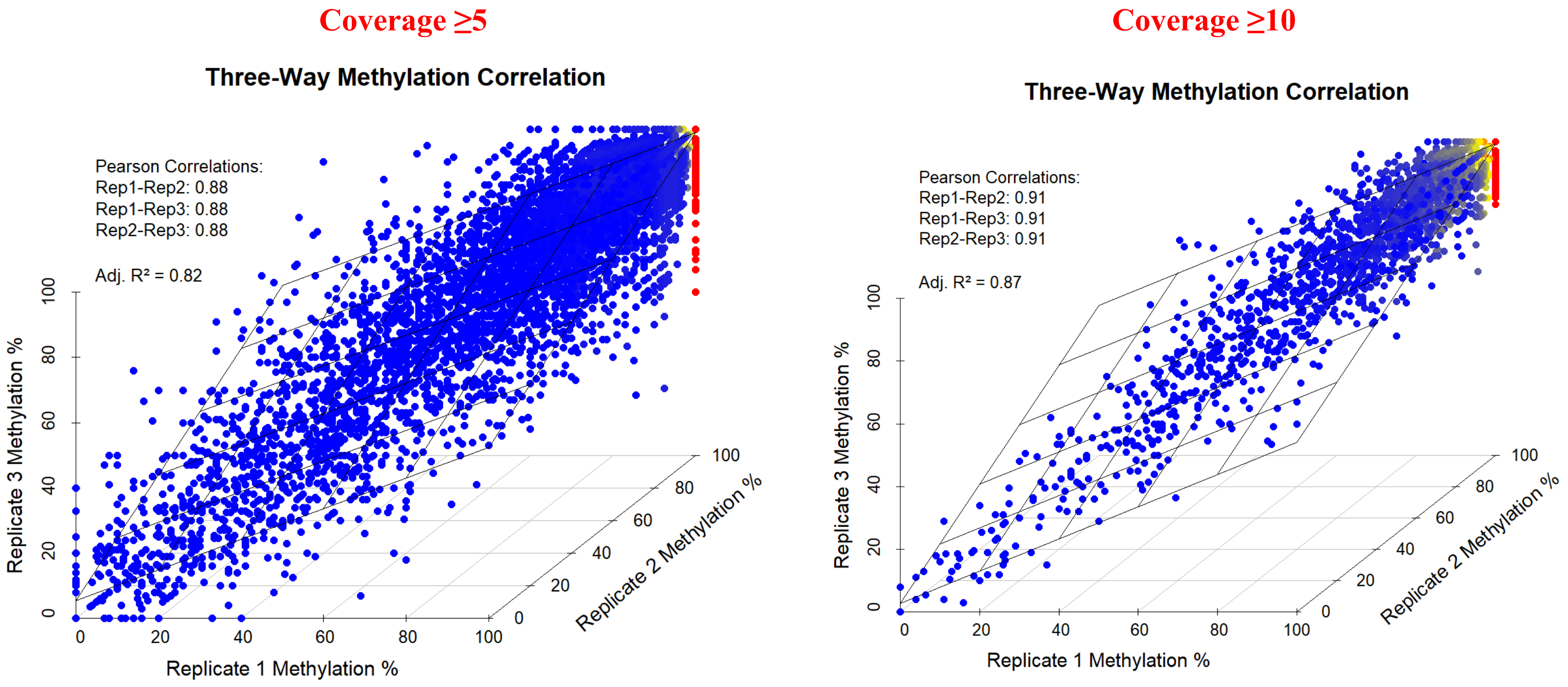


**b**

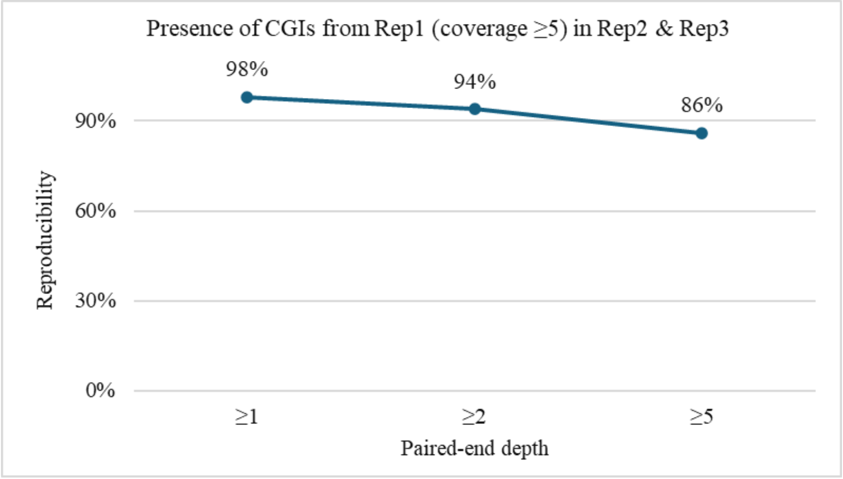


**Supplementary Fig. 2:** aMAPP reproducibility using the octamer probe GACCCGCG.

**a)** CpG methylation correlation using aMAPP on sheared gDNA from the NA12878 cell line in triplicates using coverage cut-offs of 5 and 10. **b)** Presence of covered CGIs (coverage ≥5) from Replicate 1 in Replicates 2 and 3, using aMAPP on sheared gDNA from colorectal tumor tissue.

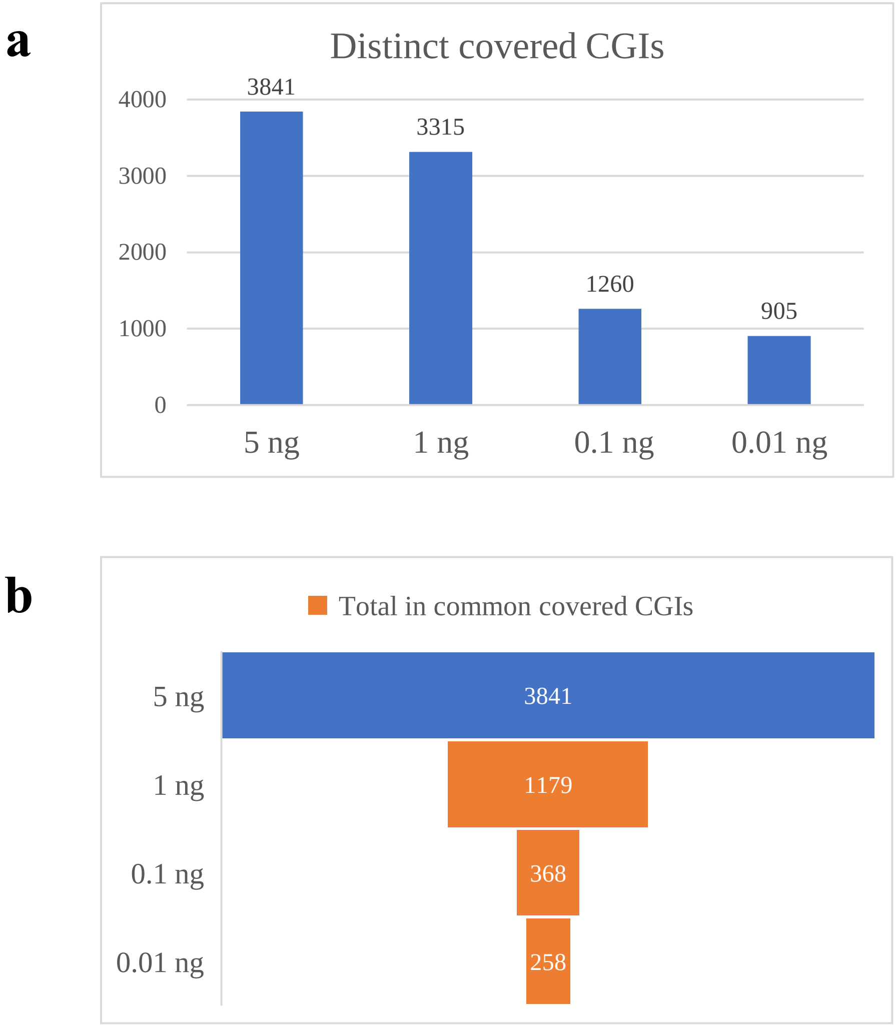


**Supplementary Fig. 3:** aMAPP capability for capturing CGIs with very low-input cfDNA using ultra-low-depth sequencing.

**a)** Number of distinct CGIs captured using aMAPP on low-input normal cfDNA. **b)** Number of commonly covered CGIs in 1ng, 0.1ng, and 0.01ng samples compared to the 5ng sample.

**a**

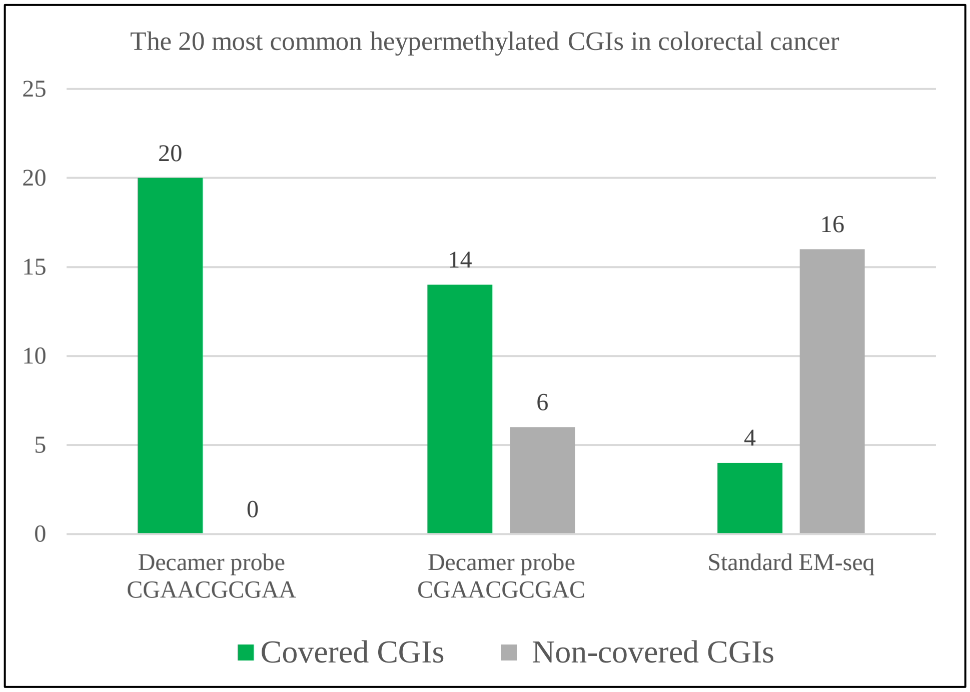


SEPTIN9_CpG 151
MLH1_CpG 93
CDKN2A_CpG 156
APC_CpG 50
MGMT_CpG 98
SFRP1_CpG 148
VIM_CpG 195
RASSF1_CpG 130
GATA5_CpG 257
GATA4_CpG 80
ICAM5_CpG 399
CDH1_CpG 103
NDRG4_CpG 192
CACNA1G_CpG 241
NEUROG1_CpG 151
IGF2_CpG 330
SOCS1_CpG 226
RUNX3_CpG 357
AXIN2_CpG 215
SDC2_CpG 160

**b**

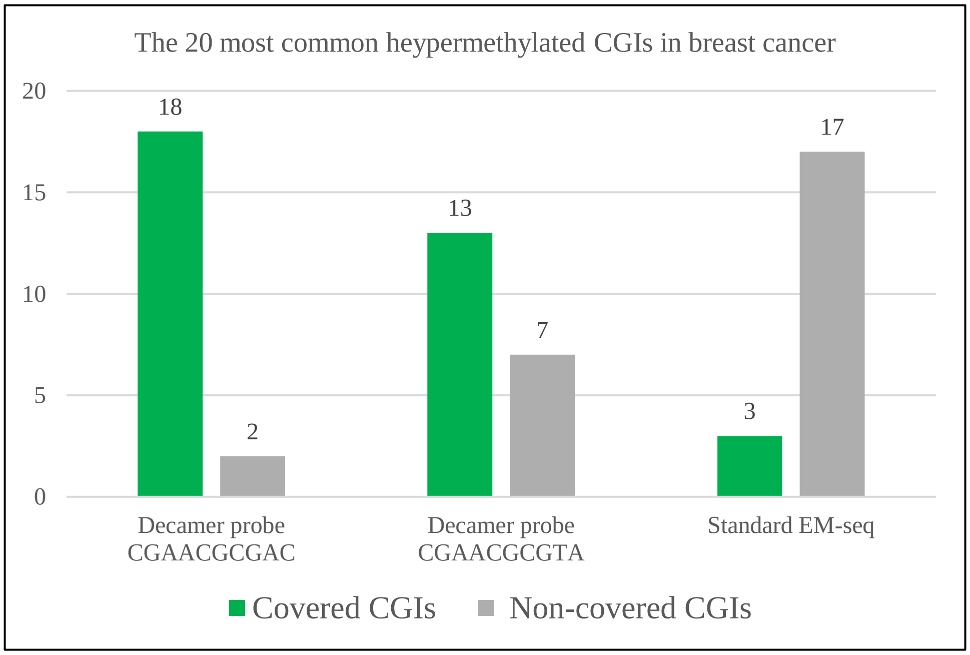


ATM_CpG 75
SOX17_CpG 215
RASSF1_CpG 130
BRCA1_promoter
GSTP1_CpG 96
HIN-1_CpG 167
APC_promoter
CCND2_CpG 379
CDH1_CpG 103
TWIST1_CpG 158
DAPK1_CpG 145
ESR1_CpG 105
CDKN2A_CpG 156
HOXA5_CpG 202
TIMP3_CpG 81
SFRP1_CpG 148
SLIT2_CpG 284
MGMT_CpG 98
GATA3_CpG 504
FOXC1_CpG 584

**Supplementary Fig. 4:** Comparison of aMAPP with different probes versus standard EM-seq in detecting the 20 most common hypermethylated CpG islands (CGIs) in colorectal and breast cancer using ultra-low depth sequencing.

**a)** Number of CGIs covered out of the 20 most common hypermethylated CGIs in colorectal cancer when aMAPP was applied using two different decamer probes, compared to standard EM-seq. **b)** Number of CGIs covered out of the 20 most common hypermethylated CGIs in breast cancer when aMAPP was applied using two different decamer probes, compared to standard EM-seq.

**a** **Methylation enrichment**

| aMAPP on EM-seq converted libraries | | Samples | | | |
|---|---|---|---|---|---|
| | | 100% Methylated | 10% Methylated | 1% Methylated | 100% Non-methylated |
| Percentage of Methylated CpGs | Genome wide | 98% | 75% | 70% | 36% |
| | RASSF1 | 100% | 100% | 82% | 11% |

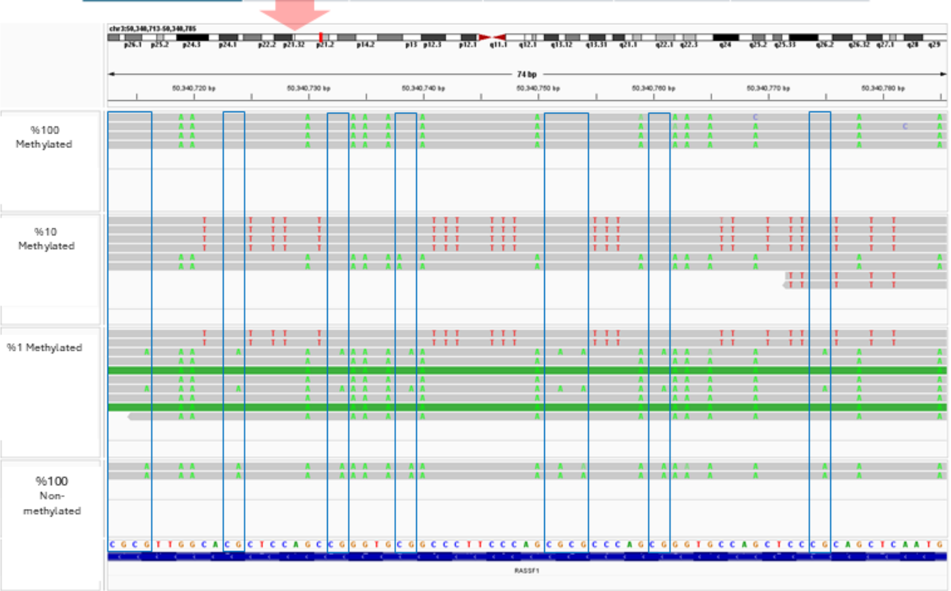


**b** **Non-methylation enrichment**

| aMAPP on TAPS converted libraries | | Samples | | | |
|---|---|---|---|---|---|
| | | 100% Non-methylated | 10% Non-methylated | 1% Non-methylated | 100% Methylated |
| Percentage of Non-methylated CpGs | Genome wide | 99% | 75% | 51% | 26% |
| | RASSF1 | 100% | 100% | 69% | No amplification |

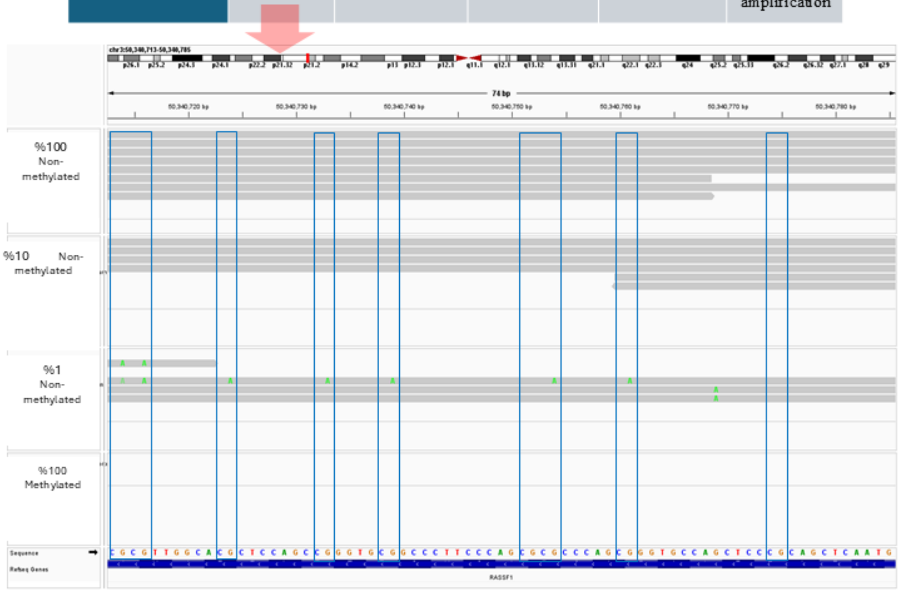


**Supplementary Fig. 5:** aMAPP's ability to trace both methylated and non-methylated CpGs at low frequencies. The enrichment effect of aMAPP was assessed using serial dilutions of control methylated DNA into non-methylated human gDNA, and vice versa, followed by ultra-low-depth sequencing. CpG methylation percentages were analyzed genome-wide and within a specific region—the CGI of RASSF1 (chr3:50,340,706–50,340,778). The methylation status of its reads was visualized using the Integrative Genomics Viewer (IGV). **a)** Methylation enrichment was observed when aMAPP was applied to EMseq-converted libraries. **b)** Non-methylation enrichment was observed when aMAPP was applied to TAPS-converted libraries.
